# Large-scale curvature sensing by epithelial monolayers depends on active cell mechanics and nuclear mechanoadaptation

**DOI:** 10.1101/2020.07.04.187468

**Authors:** Marine Luciano, Shi-Lei Xue, Winnok H. De Vos, Lorena Redondo Morata, Mathieu Surin, Frank Lafont, Edouard Hannezo, Sylvain Gabriele

## Abstract

While many tissues fold *in vivo* in a highly reproducible and robust way, epithelial folds remain difficult to reproduce *in vitro*, so that the effects and underlying mechanisms of local curvature on the epithelial tissue remains unclear. Here, we photoreticulated polyacrylamide hydrogels though an optical photomask to create corrugated hydrogels with isotropic wavy patterns, allowed us to show that concave and convex curvatures affect cellular and nuclear shape. By culturing MDCK epithelial cells at confluency on corrugated hydrogels, we showed that the substrate curvature leads to thicker epithelial zones in the valleys and thinner ones on the crest, as well as corresponding density, which can be generically explained by a simple 2D vertex model, leading us to hypothesize that curvature sensing could arise from resulting density/thickness changes. Additionally, positive and negative local curvatures lead to significant modulations of the nuclear morphology and positioning, which can also be well-explained by an extension of vertex models taking into account membrane-nucleus interactions, where thickness/density modulation generically translate into the corresponding changes in nuclear aspect ratio and position, as seen in the data. Consequently, we find that the spatial distribution of Yes associated proteins (YAP), the main transcriptional effector of the Hippo signaling pathway, is modulated in folded epithelial tissues according to the resulting thickness modulation, an effect that disappears at high cell density. Finally, we showed that these deformations are also associated with changes of A-type and B-type lamin expression, significant chromatin condensation and to lower cell proliferation rate. These findings show that active cell mechanics and nuclear mechanoadaptation are key players of the mechanistic regulation of epithelial monolayers to substrate curvature, with potential application for a number of *in vivo* situations.

## Introduction

In living systems, epithelial tissues are commonly described by three-dimensional (3D) microstructures such as invaginations, folds or wavy morphologies. Curved surfaces are also observed at interfaces between tissues or at boundaries between tissues and body lumen. The geometric form and biological function of wavy epithelial tissues are inherently linked together at all scales. For instance, crypts and villi of the small intestine provide a large surface area for exchange, improving the absorbance function. Despite its wide interest, the relationship between curvature and biological function in epithelial tissues remains largely unexplored ^1^.

Despite numerous studies on the influence of cellular and subcellular-scale topography on cell fate ^2–4^, few studies have investigated the effect of curvature on collective cell behavior ^5,6^, in particular because of the technical limitations encountered to engineer soft culture substrates with curved patterns in a controlled way ^7^. It remains unclear in particular whether and how curvatures at scales much larger than cell size could be sensed biologically. Early studies conducted on glass fibers have shown that cells orient themselves along the line of minimal curvature to minimize cytoskeletal deformations ^8,9^. Glass wires were also used to uncouple the effect of out-of-plane curvature from the lateral confinement experienced during the migration of epithelial tissues ^10^. More recently, it was shown that isolate adherent cells avoid crests of ultra-smooth sinusoidal surfaces during their migration and position themselves in valleys ^11^ and that the persistence and speed of migration of single cells can be affected by substrate curvature ^12^.

Despite these efforts, our current understanding of the role of the curvature of epithelial tissues remains elusive. To answer this question, we developed well-defined soft corrugated hydrogels to investigate the response of epithelial tissues to variations of curvature. By combining *in vitro* experiments with analytical and computational vertex model, we show that local changes of monolayer thickness and cell density can be interpreted by energy minimization arising from the mechanics of apico-lateral tensions. We extended the vertex model to consider in a minimal way changes of nuclear morphology due to active tensions and cell shape, suggesting a simple mechanism via which thickness modulation couple to the experimentally observed changes in nuclear shape and positioning. We then showed that these changes triggered by curvature also lead to high nuclear/cytoplasmic YAP ratios, demonstrating a YAP-curvature sensing of epithelial tissues, which is inhibited at high cell density. Furthermore, we showed that nuclear deformations observed on corrugated matrices were associated with a modulation of lamin A/C (LMA) and B (LMB), leading to a lower expression of LMA on positive curvatures and a higher expression of LMB on negative curvatures. Finally, we demonstrate that matrix curvature can be considered as an important regulatory cue of epithelial tissues, leading to high level of chromatin compaction and a lower DNA synthesis rate in negative curvature zones, which correspond to high cell density.

## Results

### The overall cytoskeletal architecture of wavy epithelial monolayers is not affected by curvature

To study how a monolayer of epithelial cells adapt to convex (+) and concave (−) cell-scale curvatures of their matrix, we developed a photopolymerization method based on a photoinitiator (Irgacure 2959) to generate corrugated hydroxypolyacrylamide (hydroxyPAAm) hydrogels. After UV-exposure of the hydroxyPAAm solution at 360 nm through a chromium optical photomask (Fig. 1A), the polymerization was completed in transparent zones and the slow diffusion of the Irgacure photoinitiator towards non illuminated zones lead to the formation of a smooth and wavy profile (Fig. 1B). We used transparent stripes of 10 μm wide and black stripes of 10 μm or 20 μm wide to form isotropic corrugated hydrogels of 250±30 kPa with wavelengths of 20 μm (P20, Fig. 1C) and 30 μm (P30, Fig. 1D), respectively. P20 (Fig. 1E) and P30 (Fig. 1F) corrugated hydrogels were characterized by atomic force microscopy to determine their profile, curvature of concave and convex zones, amplitude and wavelengths (table 1 and Figs. 1G-H). Corrugation patterns were symmetric and wavy structures showed periodic wavelength and constant amplitudes over large areas (10×10 mm^2^).

**Table 1.** Dimensions of P20 and P30 corrugated hydrogels. Curvature (1/R), amplitude and wavelength of P20 (n=12) and 30 (n=11) corrugated hydrogels were determined on immersed hydroxy-PAAm hydrogels by atomic force microscopy (AFM) in liquid mode. Mean ± S.D.

| Hydrogel | Curvature | Amplitude ± SD ($\mu\text{m}$) | Wavelength ± SD ($\mu\text{m}$) |
| --- | --- | --- | --- |
| P20 | 0.58 (C+) 0.13 (C-) | $2.44 \pm 0.04$ | $20.56 \pm 0.96$ |
| P30 | 0.27 (C+ and C-) | $4.62 \pm 0.16$ | $30.12 \pm 0.10$ |

**Figure 1.**
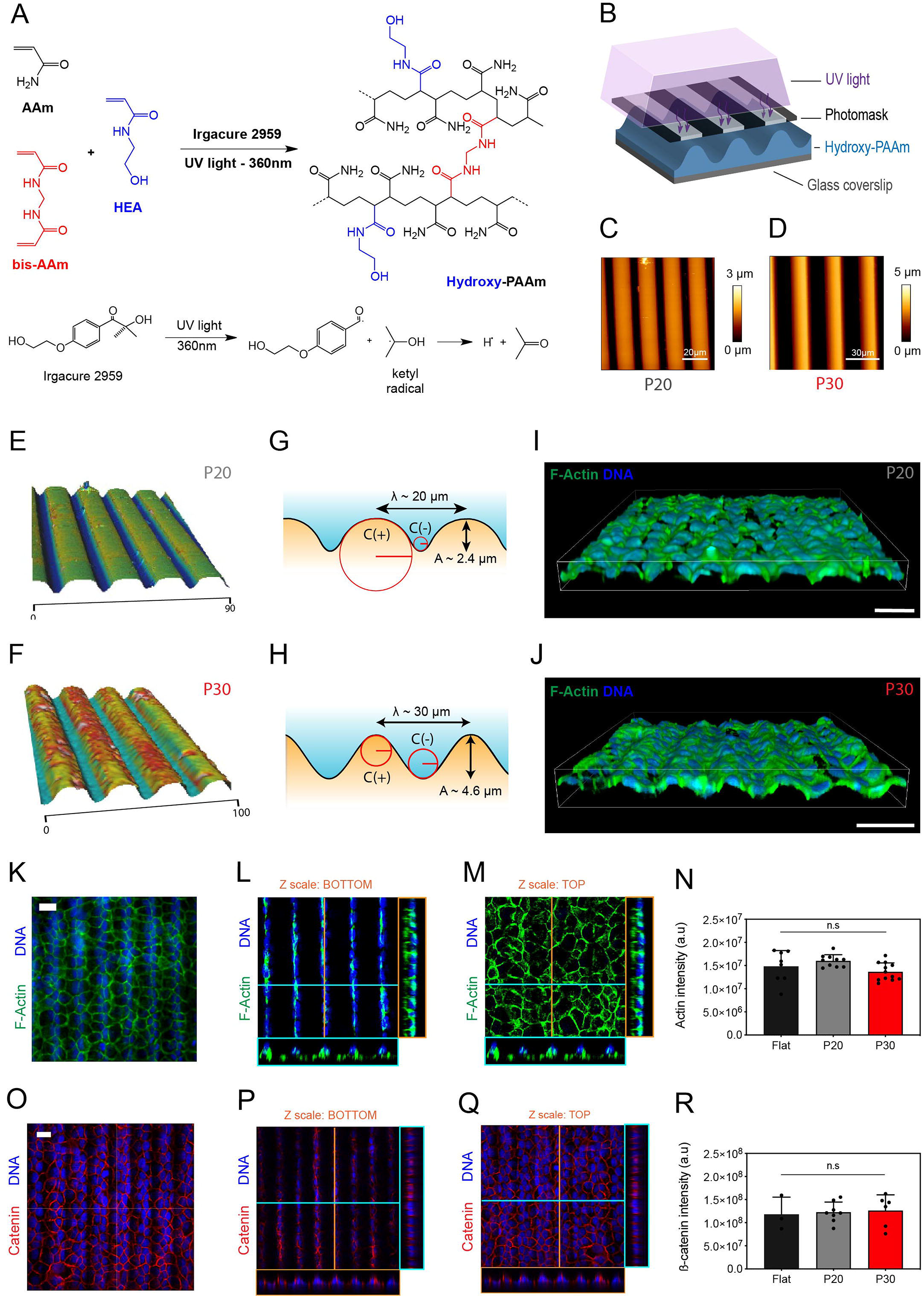
Formation of wavy epithelial monolayers on corrugated polyacrylamide hydrogels. (A) UV-Induced radical photo-polymerization of hydroxypolyacrylamide (hydroxy-PAAm) hydrogels. The radical polymerization of acrylamide (AAm, in black), bis-acrylamide (bis-acrylamide, in red) and N-hydroxyethylacrylamide (HEA, in blue) is amorced by the photoinitiator Iragure 2959 under UV exposure at 360 nm and leads to an acrylamide hydrogel with hydroxyl groups (hydroxy-PAAm). (B) Schematic representation of the UV photo-polymerization of hydroxy-PAAm hydrogels through an optical photomask. Typical atomic force microscopy (AFM) topography images of corrugated hydrogels of (C) 20 μm (P20, left) and (D) 30 μm (P30, right) wavelength. Image area is 100 μm × 100 μm for both. 3D rendering of tapping-mode topography images of (E) P20 (left) and (F) P30 (right) hydrogels. Schematic representation (side view) of (G) P20 and (H) P30 profiles with wavelengths (λ), amplitudes (A) and the radius (R) of curvature for concave (−) and convex (+) topologies. Confocal volume rendering of a MDCK epithelial monolayer grown on (I) P20 and (J) P30 corrugated hydrogels and stained for actin (in green) and DNA (in blue). (K) Maximum intensity projection and confocal orthogonal projections at (L) basal and (M) apical planes of an epithelial monolayer grown on a P30 hydrogel and stained for F-actin (in green) and nuclei (in blue). Scale bars are 30 μm. (N) Total actin intensity in epithelial tissues grown on flat (black), P20 (grey) and P30 (red) hydrogels. n=8 (flat in black), n=10 (P20 in grey) and n=12 (P30 in red). (O) Maximum intensity projection and confocal orthogonal projections at (P) the basal and (Q) the apical planes of an epithelial tissue grown on a P30 hydrogel and stained for β-catenin (red) and nucleus (blue). Total ß-catenin intensity in epithelial tissues grown on flat (black), P20 (grey) and P30 (red) hydrogels. n=10 (flat in black), n=9 (P20 in grey) and n=13 (P30 in red). n.s. is not significant.

P20 and P30 hydrogels were functionalized with human fibronectin (FN) and Madin Darby Canine Kidney (MDCK) cells were cultured at confluency (10^4^ cells/mm^2^) on flat (Supplementary Fig. S1), P20 (Fig. 1I) and P30 (Fig. 1J) corrugated hydrogels for studying how concave and convex cell-scale curvatures can affect epithelial monolayers. Epithelial tissues grown on corrugated hydrogels formed wavy confluent monolayers (Figs. 1I-J). After 48 hours in culture, wavy epithelial monolayers were immunostained with Alexafluor 488 for F-actin, 4′,6-diamidino-2-phenylindole (DAPI) for the nucleus and imaged with a laser-scanning confocal microscope. Using Z-stack projections from confocal scanning of F-actin (Figs. 1K-M, Supplementary Movie S1 and S2), we found that the actin intensity was not significantly different between flat, P20 and P30 hydrogels (Fig. 2N), suggesting that corrugations of the matrix do not affect the global amount of F-actin. We next studied the organization of cell-cell adhesive interactions in folded epithelial tissues by staining ß-catenin, which is involved in regulation and coordination of MDCK cell-cell adhesions ^13^. ß-catenin staining showed that epithelial tissues remained cohesive on corrugated hydrogels (Fig. 1O) with well-defined cell-cell contacts between polyhedral cells, independently of local concave or convex curvatures, as shown on bottom (Fig. 1P) and top (Fig. 1Q) zones, respectively. Furthermore, the global amount of ß-catenin was not significantly different between flat, P20, P30 epithelial tissues (Fig. 1R). In addition, we transformed ß-catenin stained images into skeleton images (Supplementary Figure S2A-B) to determine a mean cell area of 209.2 ± 34.3 μm^2^ on flat, 196.7 ± 27.7 μm^2^ on P20 and 182 ± 19.7 μm^2^ on P30 (Supplementary Fig. S2C), whereas the distribution of polygon classes was characteristic of the typical honeycomb pattern with 5.6 ± 1.7 on flat, 5.9 ± 1.3 on P20 and 5.8 ± 1.3 on P30 (Supplementary Fig. S2D) and characterized by an increase of the mean cell area with the polygon class, independently of the corrugations (Supplementary Fig. S2E).

**Figure 2.**
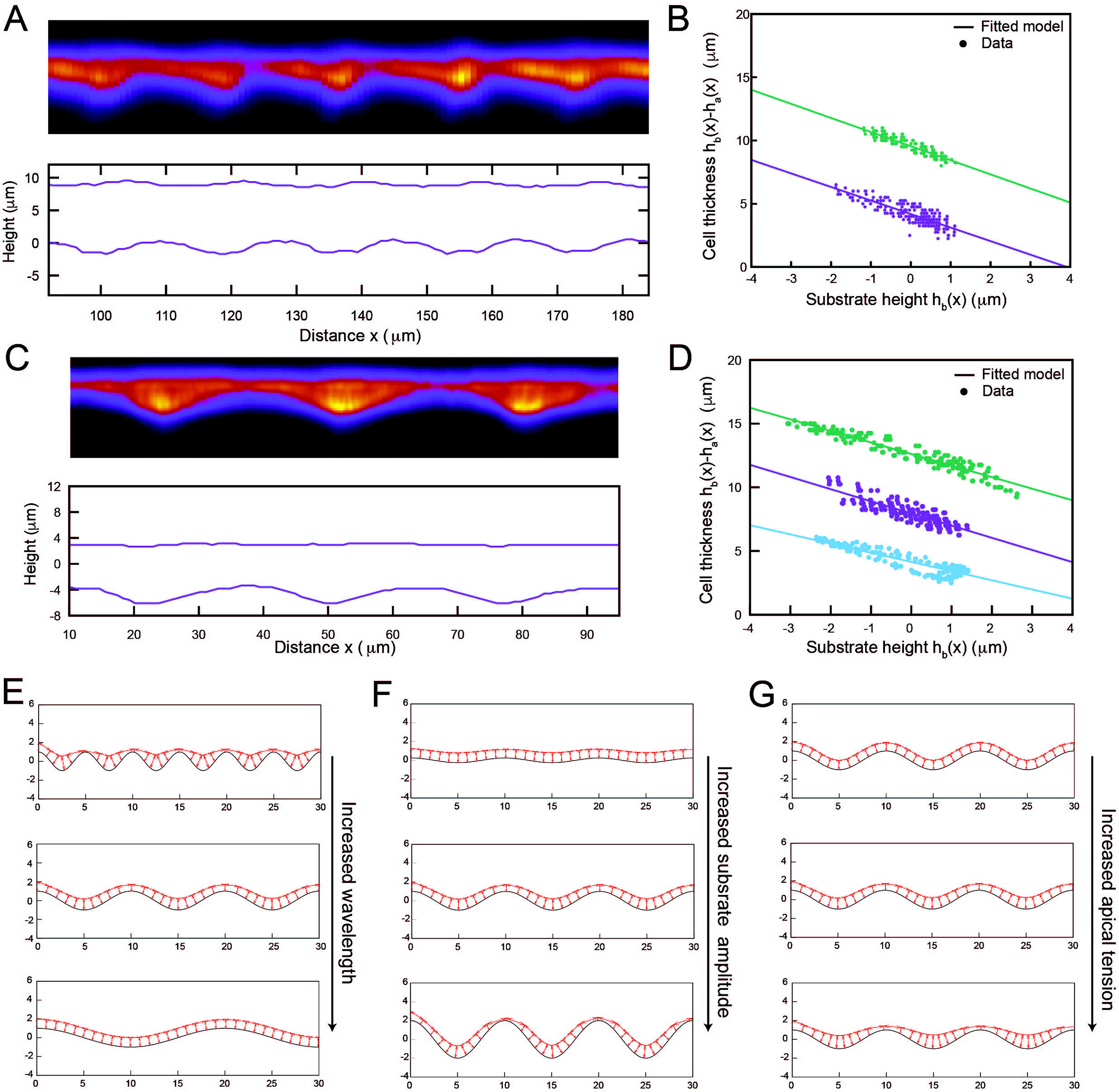
Theoretical modelling of thickness modulations from substrate curvature. (A-D) Mean-intensity projection along the pattern direction of MDCK monolayers on curved substrates (A: P20, C: P30, F-Actin staining imaged via confocal, top), together with segmented apical and basal surfaces (A: P20, C: P30). We also show cell thickness at given position x (calculated by subtracting apical and basal position in panel A,C) as a function of substrate height at the same position x (points), showing a robust negative correlation across different samples for both P20 (B) and P30 (D). Lines: theoretical prediction with fitted slope Ω = −1 ± 0.1 for P20 and Ω = −0.86 ± 0.12 for P30 (mean±S.D.), indicative of strong thickness modulation. (E-G) Equilibrium configuration of a 2D vertex model representing apical, lateral and basal surfaces of an epithelial monolayer attached to a curved substrate. (E) Increasing substrate wavelength, (F) decreasing substrate amplitude or (G) increasing apical tensions decreased thickness modulations.

Altogether, these results demonstrate that wavy epithelial monolayers maintain intact their overall architecture and can adapt to corrugated matrices without changing their hexagonal array and the global expression of actin and ß-catenin.

### Cell shape and thickness modulations from substrate curvature

However, when examining the shape of the cell monolayer as a function of curvature (Figs. 2A and 2C), we found striking differences between the top of the concave zones (i.e. crests) and the bottom of the convex zones (i.e. valleys). In both P20 and P30 tissues, cells on crests displayed reduced cell height and increased cell area (squamous-like), while cells in valleys were thicker and denser (columnar-like). We quantified this by projecting tissues along the direction of the pattern (Fig. 2A-D), thus building average intensity profiles in response to a curved substrate in each sample. We found an average 20% (resp. 40%) difference between thickness in valleys and crests in P20 (resp. P30). Interestingly, similar density differences were observed in keratinocytes ^3^, arguing this could be a general response to curvature. Given that a number of mechano-sensitive cellular responses, such as YAP localization, depend on cell thickness/density ^14,15^ as well as curvature ^2^, we thus hypothesized that curvature could be sensed indirectly by creating thickness/density differences, and turned to a theoretical model to test how curvature could create such cell shape modulations.

For this, we used the framework of vertex model ^16–19^, which describes the shape of epithelial monolayers based on their apical, lateral and basal tension ^20,21^. We first reduced the problem to 2D by neglecting the in-plane component that is translationally invariant, and assumed that the basal area could not detach from the substrate ^22^ (of amplitude *β*cos (*qx*) where *β* is the magnitude of the deformation and *q* the wavevector), thus removing its contribution to the energy. Importantly, we found that the response of the monolayer to substrate curvature depended on a single rescaled parameter, the ratio of apical to lateral tensions Γ_a_ /Γ_1_. Indeed, when apical tensions dominate, the configuration with minimum energy is the one with flat apical surface (leading to maximal thickness modulation, equal to substrate height modulation). However, this “flat-apical” configuration increases drastically total lateral area and is thus unfavorable for large lateral tensions. In that converse case, the epithelium tends to be of constant height, independent of curvature. In general, denoting acos (*qx*) the equation for the apical surface, the rescaled thickness modulation Ω = (α - β)/β is thus predicted to follow the simple law:

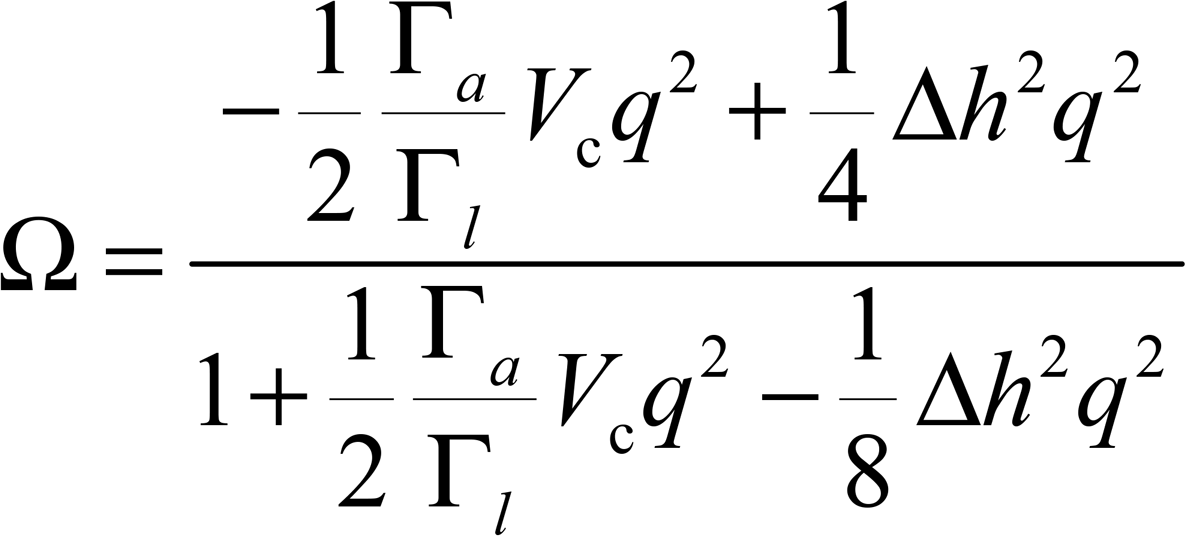

which interestingly is independent of substrate amplitude *β* (*V*_*C*_ is the cell volume and *Δh* the average thickness, see SI Text for details). We checked this analytical theory via numerical simulations by varying multiple parameters such as wavelength, amplitude, tensions, and found it matched well at large wavelength, with small-wavelength corrections (Fig. 2E-G and Supplementary Fig. S3A-C). Importantly, plotting local thickness vs. local substrate curvature in the data resulted in a linear curve (Figs. 2B,D), as predicted in the model, with the fitted slope allowing us to extract Ω (see Si Text). This value of thickness modulation was consistent with a single fitting parameter Γ_a_/Γ_1_=1-2 for P20 and P30 (other parameters such as monolayer seeding density and cell volume being constrained from independent experiments, see SI Text for details of fits and measurements). The theory thus suggests that active contractile apico-lateral tensions could be a very simple and purely physical mechanism converting changes in substrate curvature into changes in cellular thickness/densities.

### The substrate curvature modulates the morphology, distribution and orientation of the nuclei

Next, we asked whether substrate curvature would also modulate nuclear mechanics, in addition to cellular shape. We thus acquired the 3D nuclear morphologies by confocal laser scanning microscopy and found three main nuclear shapes (sketched on Fig. 3A, and shown in Supplementary Movies S3-S5), localized at the top of the concave zones (i.e. the crest), the side between concave and convex areas and the bottom of the convex zones (i.e. the valleys). As shown in Fig. 3B, nuclei adopted an oblate-shaped morphology on top of the crest, a prolate-shaped morphology in the valleys and an asymmetric shape for those accumulated on the side between concave and convex zones.

**Figure 3.**
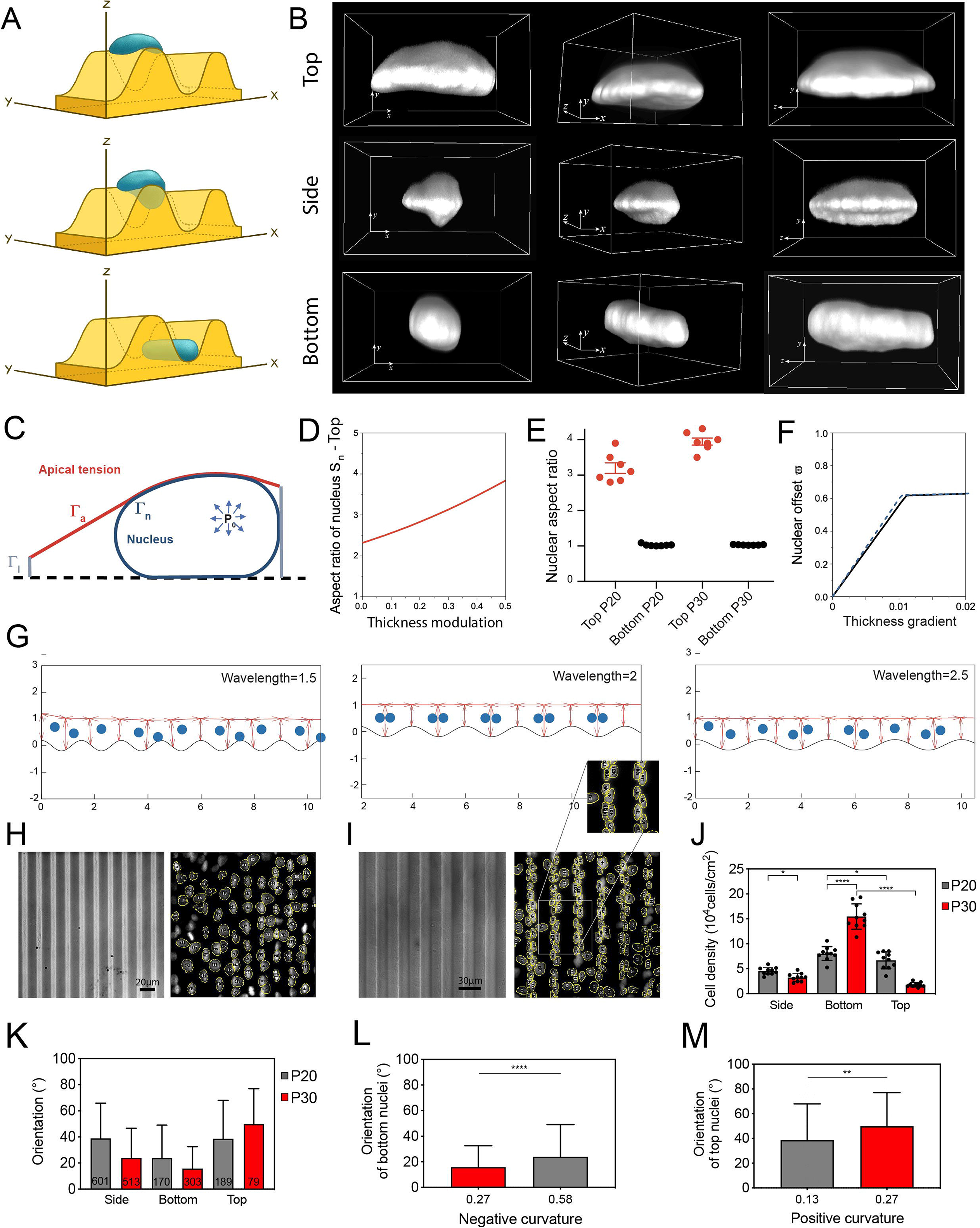
The substrate curvature modulates the morphology, distribution and orientation of the nuclei. (A) Schematic representation of the typical nuclear morphologies observed in the three main zones (top, side and bottom) of the wavy hydrogels. (B) Confocal 3D volume rendering of the typical nuclear morphologies found on the crest (top, positive curvature), on the side zone (side) and in the valley (bottom, negative curvature). (C) Sketch of the theoretical model of the cell nucleus as a tensile membrane deformed by cellular tensions, and where nucleus shape and position were calculated from force balance and depended on cell thickness and cell thickness modulation respectively. (D) Theoretical nuclear aspect ratio on crests as a function of rescaled thickness modulations. (E) Experimental nuclear aspect ratio on top/bottom of patterns (resp. red, black) for P20 and P30. (F) Theoretical nuclear positioning relative to the center of the cell as a function of thickness gradients, tending to attract cells from sides to valleys. (G) 2D vertex models (see SI Note) incorporating nuclear positioning (force proportional to thickness gradients as shown in F, taking cells away from equilibrium center position), showing that nuclei of predicted to be positioned towards valleys, an effect particularly accentuated for wavelengths at integer values of cell width (center panel). (H,I) Typical DIC images of (H) P20 and (I) P30 wavy hydrogels with the automatic detection (in yellow) of the nuclei stained with DAPI. Scale bars are 20 μm. (J) Cellular density on side, top (crest) and bottom (valleys) zones for P20 and P30 epithelial monolayer. (K) Nuclear orientation on top, side and bottom zones of P20 (in grey) and P30 (in red) wavy hydrogels. Mean orientation of the nuclei on (L) bottom (negative curvature) zones and (M) top (positive curvature) zones of P20 (in grey) and P30 (in red) wavy hydrogels. *p < 0.05, **p < 0.01, ****p < 0.0001.

We sought to test whether these features could be explained via simple first-principle physical arguments, and thus extended our vertex model to consider the interaction between a nucleus (modelled as a deformable sphere under tension) and the interfacial mechanics of a cell (Fig. 3C, Supplementary Fig. S3D for sketch, see SI Text for details). Intuitively, peaks (where cell thickness is low) are expected to compress nuclei in-plane, whereas valleys (where cell thickness is high) are expected to compress nuclei along the apico-basal axis (Fig. 3D, Supplementary Figs. S2E-H see also SI Note and Table 1 for analytical expressions). Turning to the quantitative data, we found a 3-fold difference between nuclear aspect ratio in crest vs valleys (Fig. 3E), which could be explained by the model given the previous changes in cell aspect ratio (Fig. 3D, see SI Text for details).

Furthermore, in the presence of a cell with thickness gradients /modulation (which arises generically from energy minimization on curved substrate, as detailed above), nuclei are expected to move along the thickness gradient towards high thicknesses to minimize nuclear deformation, and we predicted a linear relationship between thickness gradient and nuclear-applied deflection forces (Figs. 3F-G, Supplementary Figs. S2 D,I-J and SI Note). Importantly, these minimal theoretical considerations explained well our finding of preferential localization of nuclei towards valleys (Fig. 3G). Beyond such qualitative comparisons, the model explained highly non-trivial features of the data, including a quantization of the nuclear response with substrate wavelength when the substrate wavelength is represents integer values of cell lengths (Fig. 3G-J). For instance, for 2 cells per wavelength (close to the P30 scenario, Figs. 3G,I-J), the energy minimum of the vertex model is to have each cell spanning half-a-period between one valley and one peak, maximizing the thickness gradient and allowing full localization of nuclei in valleys. This cannot occur for non-integer values (Fig. 3G), closely matching the response to P20 substrates, where nuclei in valleys have similar shapes as nuclei in P30 valleys, but are much less frequently there (Fig. 3H-J).

Finally, concentrating now on the third dimension, along the corrugations, we found that the corrugation dimensions also modulated the orientation of the nuclei (Fig. 3K-M). Interestingly, our results showed that the high density of nuclei accumulated in the convex curvature zones (i.e. valleys) were significantly more aligned with the corrugation axis on smaller convex curvatures with 15.8±6.2° on P30 (Fig. 3L), whereas nuclei formed wider angles on large concave curvatures (i.e. crest) with 49.9±17.1° on P30 (Fig. 3M). Altogether, our findings suggest that substrate curvature has a number of feedbacks on cell and nuclear shape, and we explore in the following the accompanying biochemical changes from curvotaxis.

### Yap-curvature sensing is inhibited at high cell density

Recent works have demonstrated that direct application of forces to the nucleus was sufficient to regulate YAP activity, which is a central regulator of cell proliferation and fate, by regulating its transport through nuclear pores ^23–25^, as well as a number of nuclear mechano-transduction events being increasingly recognized ^26,27^. Given the nuclear changes we observed, this begged us to take a closer look at how YAP localization is affected by curvature. Interestingly, YAP is well-established to adapt its nuclear-cytoplasmic localization as a function of cellular density ^15^, but has also been shown to be dependent on substrate curvature via an unknown mechanism ^3^. Given our findings that curvature generically translates into density changes, we thus explored the minimal hypothesis that the latter could be a consequence of the former.

To examine the impact of substrate corrugations on YAP translocation, epithelial were grown on flat, P20 and P30 wavy hydrogels and immunolabeled after 48h for YAP and DNA (Fig. 4A). We performed high magnification confocal imaging to quantify the nuclear to cytoplasmic YAP ratio in epithelial cells located on side, bottom and top zones. We found that side and top zones of P20 (Fig. 4B) and P30 (Fig. 4C) – where cell density/thickness was low – were characterized by high nuclear/cytoplasmic YAP ratios compared to epithelial cells on flat hydrogels. However, the nuclear/cytoplasmic YAP ratio was similar on flat and bottom zones of P20 and P30. Our findings indicated therefore that the nuclear/cytoplasmic YAP ratio was larger on positive curvature zones, as predicted from the decreased cellular density there (Fig. 4D). Our results are thus consistent with the hypothesis that YAP curvature-sensing could occur indirectly from density-sensing. However, a key prediction of the theory is then that high densities should abrogate thickness modulations (as it increases the contribution of lateral tensions over apical tensions, see SI Note and Supplementary Fig. S3A), and thus YAP curvature-sensing.

**Figure 4.**
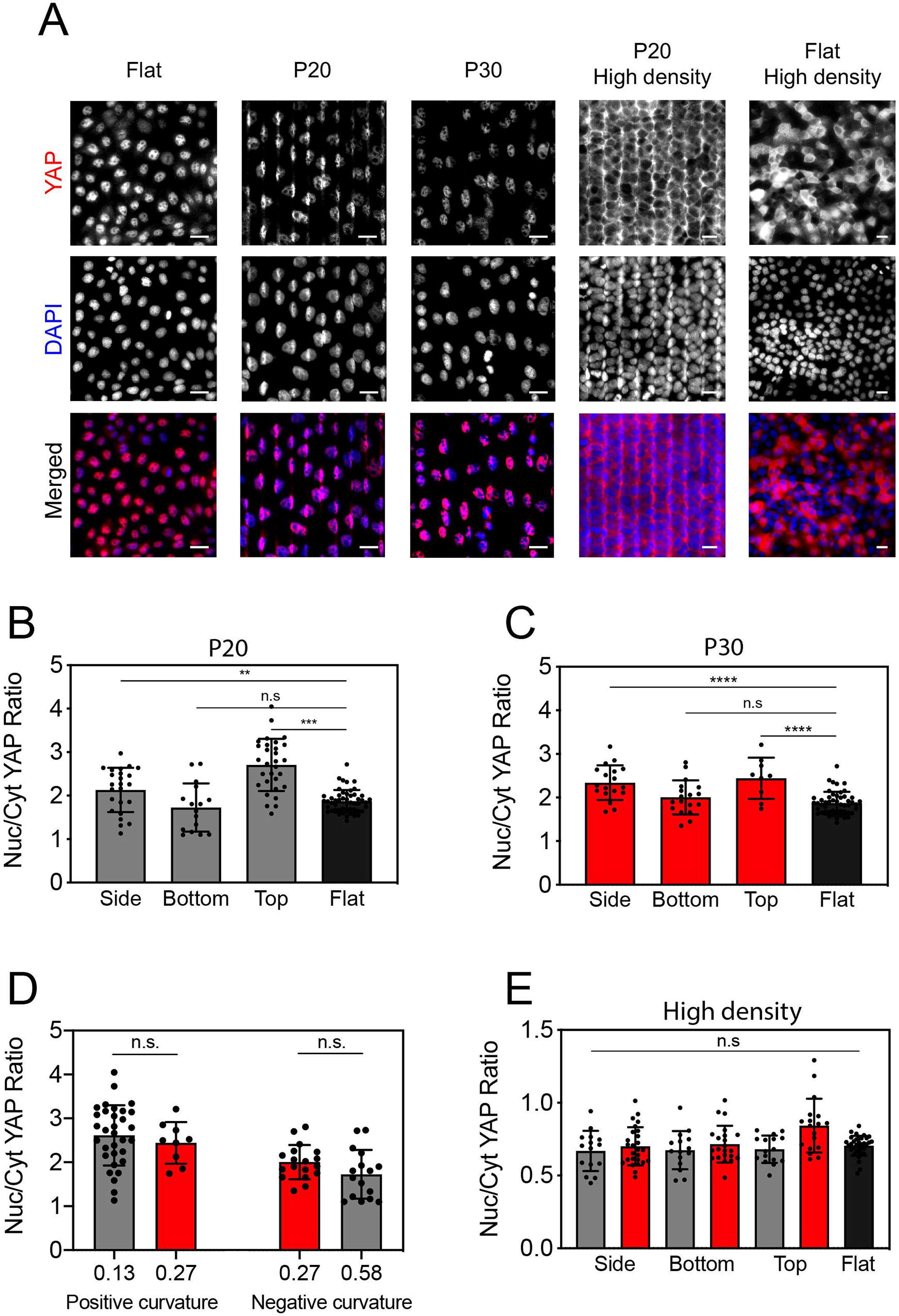
YAP-curvature sensing is inhibited at high cell densities. (A) From left to right: fluorescent images of confluent epithelial monolayers (5.10^4^ cells/cm^2^) on flat, P20 and P30 hydrogels and at high cell density (5.10^5^ cells/cm^2^) on P20 and flat hydrogels. Tissues were stained for YAP (in red) and DNA (in blue) after 48 hours in culture. Scale bars are 30 μm. Nuclear to cytoplasmic YAP ratio of nuclei on side, bottom and top zones of (B) P20 and (C) P30 corrugated hydrogels. Black bars correspond to flat hydrogels. n=30 (side), 16 (bottom), 24 (top) and 50 (flat) obtained from 5 to 7 replicates for P20 and n=18 (side), 18 (bottom), 9 (top) and 50 (flat) obtained from 5 to 9 replicates for P30. (D) Nuclear to cytoplasmic YAP ratio of nuclei in confluent epithelial monolayer in response to positive and negative curvatures. Grey bars are P20 and red bars are P30. (E) Nuclear to cytoplasmic YAP ratio of nuclei on side, bottom and top zones in confluent epithelial tissues at high cell density (5.10^5^ cells/cm^2^) on P20 (gray), P30 (in red) and flat (in black) hydrogels. n=15 (side), 15 (bottom), 12 (top) and 36 (flat) obtained from 4 to 6 replicates for P20 and n= 26 (side), 22 (bottom), 18 (top) and 36 (flat) obtained from 6 to 8 replicates for P30. All data are shown as mean ± SD. **p < 0.01, ****p < 0.0001 and n.s. not significant.

To test this prediction, we also studied very dense epithelial tissues (5.10^5^ cells/cm^2^) on P20 and flat hydrogels ^28,29^, and found a low nuclear/cytoplasmic YAP ratio, which was unaffected by substrate curvature (Fig. 4E), confirming that high cell density can inhibit YAP curvature-sensing from epithelial cells. Finally, we sought to apply our theory to previously published dataset on keratocytes, and found that we could again explain multiple features of these within our theory (with larger apical/lateral tension ratio Γa/Γ1=5, see SI Text and Fig. S3), including the magnitude of valley/crest density differences, and the delamination of cells from the crests above a critical value of substrate curvature (see Supplementary Fig. S3B and SI Text for details). This argues for a generality of such a mechanism of density/thickness changes from curvature, with impact on YAP.

### The substrate curvature modulates the relative abundance of nuclear lamina

Given our findings that matrix curvature leads to differential nuclear deformations, we sought to evaluate the lamina composition in deformed nuclei, as a potential mechano-sensitive response ^30^. Indeed, accumulating evidence shows that the nuclear mechanical response can be characterized as a combination of elastic (spring-like) and viscous (liquid-like) properties ^31^, described by the stoichiometric ratio between B-type (LMB) and A-type (LMA) lamins ^32^. In this model, B-type lamins contribute primarily to the elastic response, whereas A-type lamins contribute to the viscosity ^33,34^. We examined by scanning confocal microscopy the fluorescence intensity of A-type (TRITC in red) and B-type (FITC in green) lamins for the three main nuclear morphologies (bottom, side and top) on P20 corrugated hydrogels (Fig. 5A-D). Z-stack images using a ×60 objective were collected for three channels (DAPI, TRITC and FITC) from the entire volume of the nuclei using a step size of 0.15 μm. Exposure times and laser power were kept constant and acquired stack of images was deconvolved to remove out of focus light.

**Figure 5.**
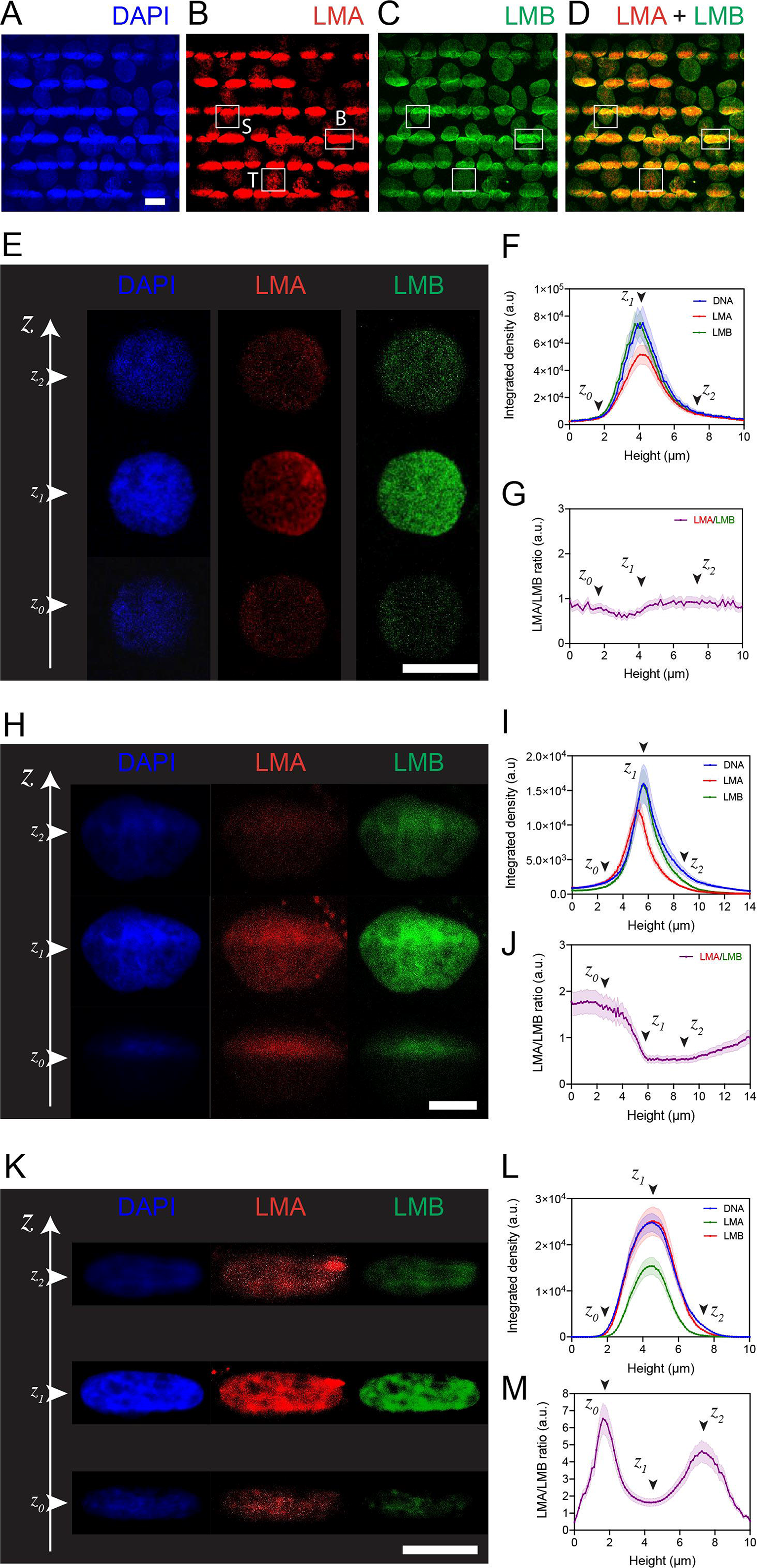
The nuclear lamina is responsive to the substrate curvature. Maximum intensity projection images of the nuclei of MDCK epithelial monolayer grown on a P20 hydrogel and stained with (A) DAPI in blue, (B) LMA in red and (C) LMB in green. (D) Merge image of LMA and LMNB. The scale bar is 20 μm. The white rectangles correspond to nuclei located on top (T), side (S) and bottom (B) zones of the corrugated hydrogel. Confocal Z stack images of nuclei stained with DAPI in blue, LMA in red and LMB in green and located (E) on the top of a positive crest, (H) on the side between valleys and crest and (K) in the bottom of the valley. Z_0_, Z_1_ and Z_2_ correspond to the basal, intermediate and apical zones, respectively. Fluorescence intensity of DAPI (in blue), LMA (in red) and LMB (in green) versus nuclear height (F) on top, (I) on side and (L) in the bottom. LMA to LMB intensity ratio versus the nuclear height (F) on top, (I) on side and (L) in the bottom. Black arrows indicate Z_0_, Z_1_ and Z_2_ apical zones. All data are shown as mean ± S.D. (n=4 for each).

Using Z-stack images of nuclei located on top (Fig. 5E, Supplementary Movie S6), side (Fig. 5H, Supplementary Movie S7) and bottom (Fig. 5K, Supplementary Movie S8) zones, we estimated the mean integrated density of DAPI, LMA and LMB for each Z-plane and calculated the LMA/LMB integrated density ratio to characterize the relative abundance of A- versus B-type lamins. For each nuclear morphology, we defined three Z planes of interest based on the DAPI curve: Z_1_ (mid plane) as the maximum, *Z*_0_ (basal plane) as Z_1_-2σ and Z_2_ (apical plane) as Z_1_+2σ. The mean integrated densities of DAPI, LMA and LMB versus the nuclear height for the three nuclear morphologies (n=8 for each) were characterized by gaussian-like distributions (Figs. 5F, 5I and 5L). We observed similar distributions of LMB and DAPI in top nuclei and a lower integrated density for LMA. The LMA/LMB ratio for top nuclei was slightly below 1 for the apical part (Z_0_, Fig. 5G) and then close to 1 for Z>Z_1_, indicating that the LMA abundance was lower on positive curvatures. The distributions of LMB and DAPI in side nuclei were both located at 5.7 μm, whereas the maxima of the LMA distribution was centered at 5.2 μm suggesting a higher abundance of LMA in the basal part of side nuclei. The LMA/LMB ratio was ~1.7 at Z_0_, then decreased abruptly to Z_1_ to reach a value of ~0.5 for Z_1_<Z<Z_2_. The asymmetric morphology of side nuclei was therefore characterized by more LMA in the basal part. Interestingly, our findings showed that LMB and DNA integrated densities exhibited a similar distribution (Fig. 5L), whereas the LMA integrated density was lower. The LMA/LMB ratio indicated that the relative abundance of LMB was significantly higher at Z_0_ and Z_2_ focal planes (Fig. 5M), corresponding to the basal and apical sides of the nuclei aligned into the valleys.

Taken together, out results indicated that the nuclear deformations lead to a lower abundance of LMA on positive curvatures and a higher abundance of LMB on negative curvatures, demonstrating a modulation of the relative abundance of A- versus B-type lamins in deformed nuclei in response to the substrate curvature.

### Concave curvatures lead to lower cell proliferation rate and promote significant chromatin condensation

Finally, the modulation of nuclear shape, YAP localization and the relative lamin A/B abundance by substrate curvature led us to question how matrix corrugations affect cell functions, such as proliferation and DNA synthesis.

We first checked whether nuclear volume was affected by curvature, using high magnification confocal Z-stacks of nuclei located on side, bottom and top zones of P20 (Fig. 6A) and P30 (Fig. 6B) corrugated hydrogels. Cells on top and bottom zones of P20 and P30 hydrogels exhibited lower nuclear volumes that those on side zones, suggesting that the nuclear volume was decreased by both positive and negative curvatures. Interestingly, the lower nuclear volumes were found in cells located on the bottom zones of P20 (635 ± 233 μm^3^) and P30 (776 ± 116 μm^3^), demonstrating that the nuclear shape remodeling in negative curvature zones, which is associated high cell density and prolate-shaped nuclei, resulted in a significant nuclear volume loss.

**Figure 6.**
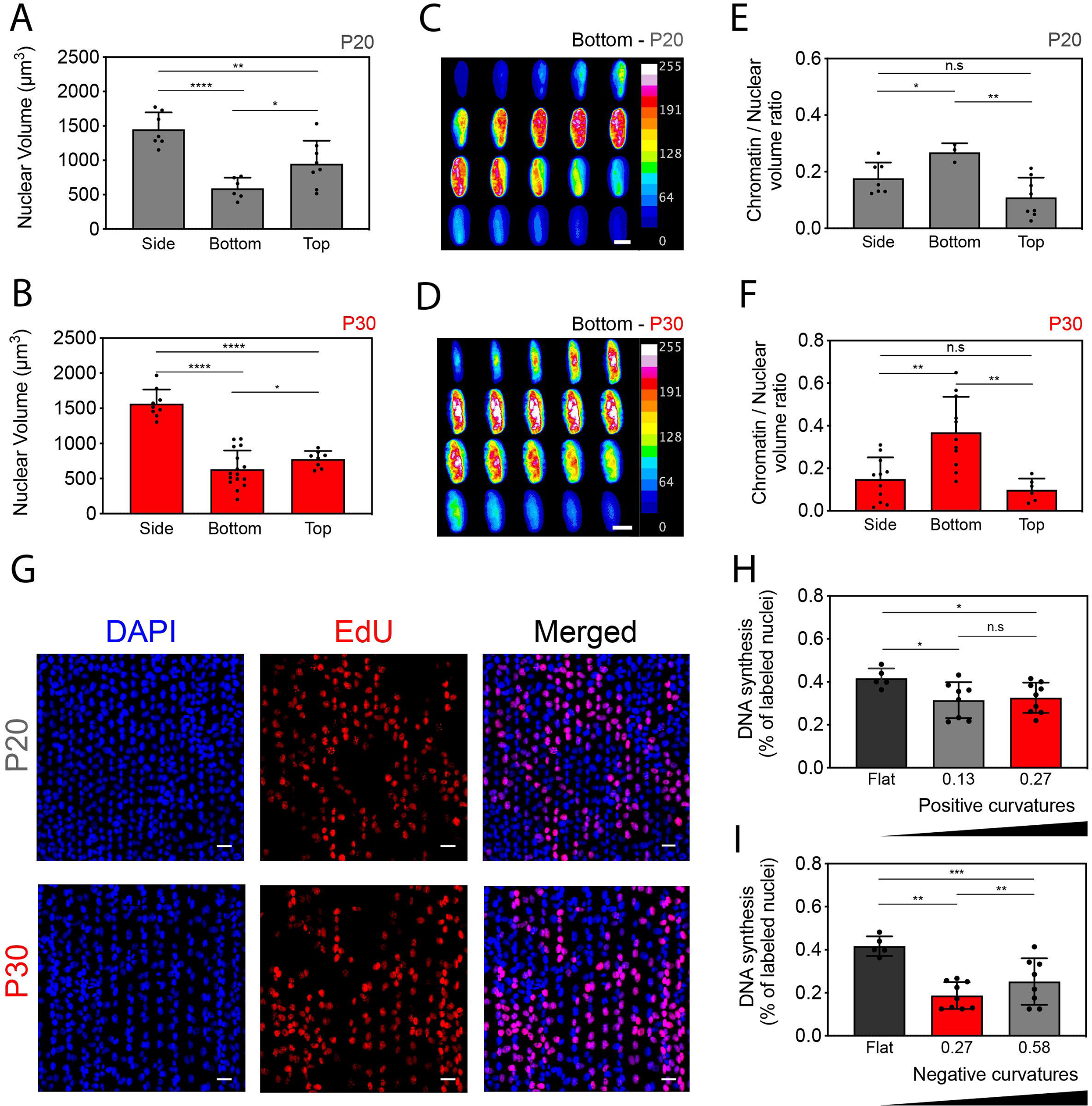
Concave curvature zones lead to lower cell proliferation rate and promote significant chromatin condensation in elongated nuclei. Nuclear volume for cells localized on side, bottom and top zones of (A) P20 (in grey) and (B) P30 (in red) corrugated hydrogels. Typical successive changes of the level of chromatin condensation in a nucleus localized in (C) a bottom zone of P20 wavy hydrogel and (D) a side zone of a P30 wavy hydrogel. Intensities of DNA staining were digitized in 256 bits and color coded for each Z-stack. Highly condensed domains show higher fluorescence intensity with respect to the less condensed ones. Chromatin to nuclear volume ratio for cells localized on side, bottom and top zones of epithelial tissues seeded on (E) P20 and (F) P30 wavy hydrogels. (G) Typical images of DAPI (in blue) and Edu (in red) staining in an epithelial tissue grown on a P20 (top row) and P30 (bottom row) corrugated hydrogels. Scale bars are 20 μm. Rate of Edu-positive cells on (H) positive and (I) negative curvatures. Gray bars are P20, red bars are P30 and black bars are flat hydrogels. All data are shown as mean ± SD. *p < 0.05, **p < 0.01, ***p < 0.001, ****p < 0.0001 and n.s. not significant.

Secondly, we checked whether substrate curvature also affected the chromatin organization, using a quantitative procedure based on DAPI staining ^35,36^. Indeed, the uptake of DAPI depends on the total amount of DNA, but also on its level of condensation ^37^. The average spatial density corresponding to the ratio between the integrated fluorescence intensity and the volume of the nucleus is therefore a reliable indicator of the *in-situ* average chromatin condensation. As shown in Figs 6C and 6D, marked reorganization of chromatin distribution was associated with nuclear deformation in bottom zones of P20 and P30 corrugated hydrogels. Highly condensed chromatin domains showed higher fluorescence intensity with respect to the less condensed ones. The quantification of the chromatin to nuclear ratio indicated that nuclei deformed in the bottom zones of P20 (Fig. 6E) and P30 (Fig. 6F) showed the highest chromatin compaction values with 0.27 ± 0.03 and 0.37 ± 0.16, respectively. Our results indicated that prolate nuclear shapes in negative curvature zones were associated with a high level of chromatin compaction that may affect DNA synthesis.

To test this hypothesis, we used the incorporation of a thymidine analog, 5-ethynyl-2′-deoxyuridine (EdU), as a proliferation marker. A 24-hour incubation period was chosen because it allowed EdU incorporation on corrugated and flat hydrogels, without no saturation of EdU incorporation in MDCK cells. The Edu-positive ratio was calculated as the ratio between Edu and Hoechst stained cells in a particular field of view (Fig. 6G). We observed lower rates of positive-EdU nuclei on positive curvature than on flat hydrogels, regardless the positive curvature value (Fig. 6H). In addition, our findings showed lower rate of positive-EdU nuclei in the valleys, demonstrating that the DNA synthesis was significantly decreased in cells located on negative curvature zones.

Taken together, these results indicated that negative curvatures zones of corrugated matrices lead to the deformation of the nuclei in a prolate shape, which is associated with a high level of chromatin compaction and a lower DNA synthesis rate.

## Discussion

The role of the substrate curvature has been mainly described so far at the single-cell level, establishing that cells orient themselves along the line of minimal curvature to minimize cytoskeletal deformations. Here, we explore the hypothesis that confluent epithelial monolayers can also sense large-scale curvature via active cell mechanics and nuclear mechano-sensing. Our findings reveal that substrate curvature leads to thicker epithelial zones in the valleys and thinner ones on the crest, as well as corresponding cellular densities. This is fully recapitulated in a physical model of apico-lateral active tension, where thickness modulation arises generically as stable states of the monolayer (and consistent with findings of a pre-print released during the preparation of this manuscript ^38^, leading us to hypothesize that matrix curvature sensing could arise from resulting density/thickness changes. Furthermore, we develop a minimal theory for how thickness modulations imposed by the substrate can impact nuclear morphometrics, such as nuclear deformation, aspect ratio and positioning relative to the local curvature. These predicted flattened nuclei at the top of the patterns, but also a movement of nuclei towards the valleys in order to minimize their deformation in response to thickness gradients. We verified these features in experiments, including non-trivial features such as enhanced responses to specific wavelengths. Together, these findings show that physical processes allow epithelial monolayers to respond to curvature changes, leading to the appearance of different types of nuclear deformations and orientations.

Given accumulating evidence of nuclear mechano-transduction processes ^27,39–41^ we then investigated how these physical changes translated into mechano-responses and biochemical changes within the cells as a function of local curvature. We showed that the patterns of cellular densities generated by the matrix curvature are associated with significant changes in the spatial distribution of YAP ^2,3^, consistent with the idea of curvature-sensing occurring via physically-driven thickness/density-changes. YAP in particular has been shown to be is a key transcription factor that mediates the interplay between cellular mechanics and signaling cascades underlying gene expression, cell proliferation, differentiation fate decisions, and organ development. The spatio-temporal localization of YAP provides therefore critical information about the regulatory state of the cell and in the future, it will be interesting to probe the generality of these findings to other cellular lines and physiological contexts.

In addition to YAP response, our results also indicated that lamin A/C ratio ^42,43^, chromatin condensation and cell proliferation rate are all modulated by substrate curvature, demonstrating that substrate curvature sensing modulates multiple mechanotransduction pathways in cell assemblies. The correlation of YAP and lamins with stem cell differentiation and cell division has suggested recently a relationship between the expression of both proteins in deformed nuclei ^44^, which would be a natural next step of investigation. Our study therefore provides insights into the mechanistic regulation of epithelial monolayers to large-scale substrate curvature, with potential impact for a large number of physiological situations, given the ubiquitous presence of curvature *in vivo.*

## Materials and Methods

### Fabrication of corrugated polyacrylamide hydrogels by UV-photocrosslinking

Instead of the standard radical polymerization using catalysts such as tetramethylenediamine (TEMED) and ammonium persulfate (APS), which lead to slow polymerization times, we used a photoinitiator (Irgacure 2959) to polymerize hydroxypolyacrylamide (hydroxy-PAAm) hydrogels. Hydroxy-polyacrylamide (hydroxy-PAAm) hydrogels were prepared by mixing acrylamide (AAm), bis-acrylamide (bis-AAm), N-hydroxyethylacrylamide (HEA), 2-Hydroxy-4′-(2-hydroxyethoxy)-2-methylpropiophenone (Irgacure 2959) and deionized water. A solution composed of 2836 μL of acrylamide (AAm, Sigma #79-06-1) at 15% w/w in deionized water, 1943 μl of N,N’- methylenebisacrylamide (BisAAm, Sigma #110-26-9) at 2% w/w in deionized water, and 1065 μL N-hydroxyethylacrylamide monomers at 65 mg/mL in deionized water (HEA, Sigma #924-42-5) were mixed together in a 15 mL Eppendorf tube ^45–47^. Deionized water was added to reach a final volume of 6 mL and then 1 ml of the Irgacure 2959 photoinitiateur (2-Hydroxy-4′-(2-hydroxyethoxy)-2- methylpropiophenone, Sigma) was added to the mixture. After a gentle mixing, the solution was degassed during 30 min under a nitrogen flow. Glass coverslips of 22 mm^2^ in diameter were cleaned with 0.1 M NaOH solution during 5 min and then rinsed abundantly with deionized water during 20 min under agitation. Cleaned glass coverslips were then treated during one hour with 3-(trimethoxysilyl)propyl acrylate (Sigma #2530-85-0) to promote a strong adhesion between the hydroxy-PAAm hydrogel and the glass coverslips and finally dried under a nitrogen flow. A volume of 40 μL of the degassed mixture was squeezed between an activated glass coverslip and a chromium optical photomask (Toppan photomask, France) and before exposition to UV illumination at 360 nm (Dymax UV light curing lamp). Chromium optical photomasks with alterning transparent stripes of 10 μm wide and black stripes of 10 μm or 20 μm wide were used to form corrugated hydrogels with wavelengths of 20 μm (P20) and 30 μm (P30), respectively. After UV exposition of 10 min at 10 mW/cm^2^ through the optical photomask, the polymerization was completed and a corrugated hydroxy-PAAm hydrogel was formed. Finally, hydrogels were gently removed from the photomask under water immersion, washed three times in sterile deionized water under gentle agitation and stored in sterile deionized water at 4°C.

### Characterization of corrugated hydrogels by atomic force microscopy

Atomic force microscopy (AFM) measurements were performed on corrugated hydrogels immersed in PBS at 7.2 pH. Once the coverslip containing the gel was placed on the stage of the microscope, the cantilever was positioned far above the glass surface and allowed to thermally equilibrate during 30 min. AFM measurements were performed using a NanoWizard TM 3 AFM (JPK Instruments, Berlin, Germany) operated in Quantitative Imaging (QI) mode. Cantilevers were purchased from Bruker (MLCT-BIO-DC-F: k = 0.6 N/m, f = 125 kHz; silicon nitride; front angle 35 ± 2°; quadratic pyramid tip shape). Cantilever spring constant and sensitivity were calibrated before each experiment using the JPK software. The force trigger was adjusted to have a high indentation without damaging the gel. Tip velocity was maximized within instrument limits and ramp size was reduced with a short baseline in order to minimize the acquisition time. Data were analyzed with the NanoWizard^®^ Data Processing software version 6.1.96 and the stiffness was calculated according the Hertzian contact model (Young’s modulus), fitting the distribution to Gaussians.

### Cell culture

Epithelial cells from the Madin-Darby Canine Kidney cell line (MDCK II, Sigma #85011435) were maintained in polystyrene T75 flasks of in a cell culture incubator at 37°C and 5% CO_2_. MDCK cells were cultured in proliferation medium composed of Dubelcco’s Modified Eagle’s medium (DMEM), high glucose (4.5 g/l) with L-glutamine (BE12-604F, Lonza) supplemented with 10% (v/v) Fetal Bovine Serum (FBS, AE Scientific) and 1% of penicillin and streptomycin antibiotics (AE Scientific) ^48^. MDCK cells were seeded on flat (control), P20 and P30 corrugated hydrogels at a density of 5.10^4^ cells/cm^2^ for 48 hours. Few experiments were performed at high density (5.10^5^ cells/cm^2^) for 48 hours.

### Proliferation assays

The S-phase synthesis of the cell cycle was labeled in living epithelial tissues grown on corrugated hydrogels using the Click-iT EdU (5-ethynyl-2’-deoxyuridine) (Thermofischer Scientific, A20174) for 30 minutes in proliferation media. Proliferation of MDCK cells on corrugated hydrogels was assessed using the Click-iT EdU Alexa (647 or 555) kit (Thermofischer Scientific, C10338). Briefly, epithelial tissues were incubated with a 10 μM solution of EdU in complete medium for 30 min. Then, tissues were rinsed with PBS, fixed for 10 min with a 4% PFA solution and permeabilized for 20 min with a 0.5% solution of Triton X-100. MDCK tissues cells were blocked with 3% BSA and incubated for 30 min in the dark with a reaction cocktail composed of Click-iT reaction buffer, CuSO4, AlexaFluor azide and reaction buffer additive. MDCK tissues were rinsed with PBS three times and labelled with Hoechst 33342 before mounting in slow fade gold antifade.

### Immunostaining of epithelial tissues

MDCK cells were fixed and permeabilized with 4% paraformaldehyde (Electron Microscopy Sciences), 0.05% Triton X-100 (Sigma) in phosphate buffered saline (PBS 1X, Capricorn scientific) for 12 min at room temperature. Fixed cells were rinsed three times in warm PBS and incubated 30 min with a blocking solution containing 1% BSA (Bovine Serum Albumine, GE Healthcare) and 5% FBS in PBS. MDCK cells were labeled for F-actin with Alexa Fluor 488 Phalloidin, 1:200, DNA with DAPI at 1:200 (ThermoFisher Scientific, #D1306) and ß-catenin ^49^. Yap was labeled with YAP1 monoclonal antibody produced in mouse (Abnova, clone 2F12), ß-catenin with anti ß-catenin produced in mouse at 1:200 Sigma-Aldrich, #C2206) for 45 min at 37°C. MDCK cells were washed three times in PBS, incubated with an anti-mouse antibody produced in goat and labeled with a goat anti-mouse antibody at 1:200 (Molecular Probes, tetramethylrhodamine, Invitrogen, T2762) for 45 min at 37°C. Immunostained cells were mounted on microscope slides with slowfade gold antifade (Thermofisher, Molecular probes) for epifluorescence and confocal imaging.

### Epifluorescence and confocal imaging

MDCK epithelial tissues were observed in epifluorescence and confocal mode with a Nikon Eclipse Ti-E motorized inverted microscope equipped with x20, x40 and x60 Plan Apo (NA 1.45, oil immersion) objectives, two lasers (Ar ion 488 nm; HeNe, 543 nm) and a modulable diode (408nm). Epifluorescence images were recorded with a Roper QuantEM:512SC EMCCD camera (Photometrics Tucson, AZ) using NIS Elements Advances Research 4.0 software (Nikon). Confocal images using small Z-depth increments (0.15 μm) were processed using NIS-Elements (Nikon, Advanced Research version 4.0).

### Morphometric analysis of the nuclei

The morphometric analysis of the nuclei was conducted using the the trainable WeKa segmentation plugin for FIJI ^50^ that first calculates the barycenter of the nuclei and then extracts contour, perimeter and projected area of each nuclei. Nuclei located at the border of the image are automatically rejected from the working population. Then the algorithm generates an ellipse to fit the nuclear contour and determine the orientation angle of each nuclei with respect to the orientation of the corrugations.

### Statistical analysis

Differences in means between groups were evaluated by two-tailed Student’s t-tests performed in Prism 7b (Graphpad Software, Inc.). For multiple comparisons the differences were determined by using an analysis of variance (ANOVA) followed by Tukey post-hoc test. *p < 0.05, **p < 0.01, ***p < 0.001, ****p < 0.0001 and n.s. not significant. Unless otherwise stated, all data are presented as mean ± standard deviation (S.D.).

## Acknowledgments

This project was supported by the European Research Council under the European Union’s Horizon 2020 Research and Innovation Program Grant Agreements 851288 (to E.H), and by the Austrian Science Fund (FWF) (P 31639 to E.H.). L.R.-M. acknowledges funding from the Agence National de la Recherche (ANR), as part of the “Investments d’Avenir” Programme (I-SITE ULNE / ANR-16-IDEX-0004 ULNE). This work benefited from ANR-10-EQPX-04-01 and FEDER 12001407 grants to F.L. This work was conducted with the financial support from FEDER Prostem research project no. 1510614 (Wallonia DG06) to S.G. M.L. is financially supported by FRIA (F.R.S.-FNRS). M.S. is senior research associate of the Fund for Scientific Research (F.R.S.-FNRS) and acknowledges the grant EOS No. 30650939 (PRECISION). S.G. acknowledges the Interreg MAT(T)ISSE project, which is financially supported by Interreg France-Wallonie-Vlaanderen (Fonds Européen de Développement Régional, FEDER-ERDF). Winnok H De Vos acknowledges FWO Grant number G005819N.

