## Supplementary Information for "Large-scale curvature sensing by epithelial monolayers depends on active cell mechanics and nuclear mechanoadaptation"

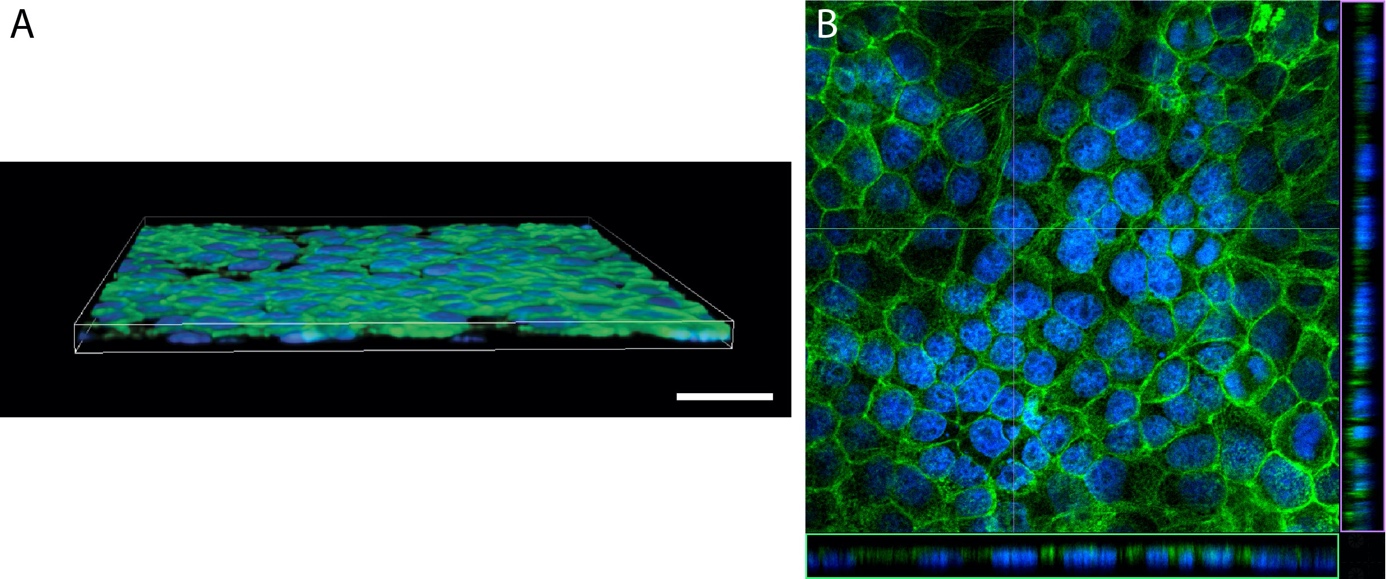

**Supplementary Figure S1 -** **Epithelial monolayer on flat hydrogels.** (A) 3D volume rendering and (B) orthogonal view of a MDCK monolayer grown on a flat hydroxy-PAAm hydrogel coated with FN. Actin is labeled in green with AlexaFluor 488 and DNA in blue with DAPI. The scale bar is 55 µm.

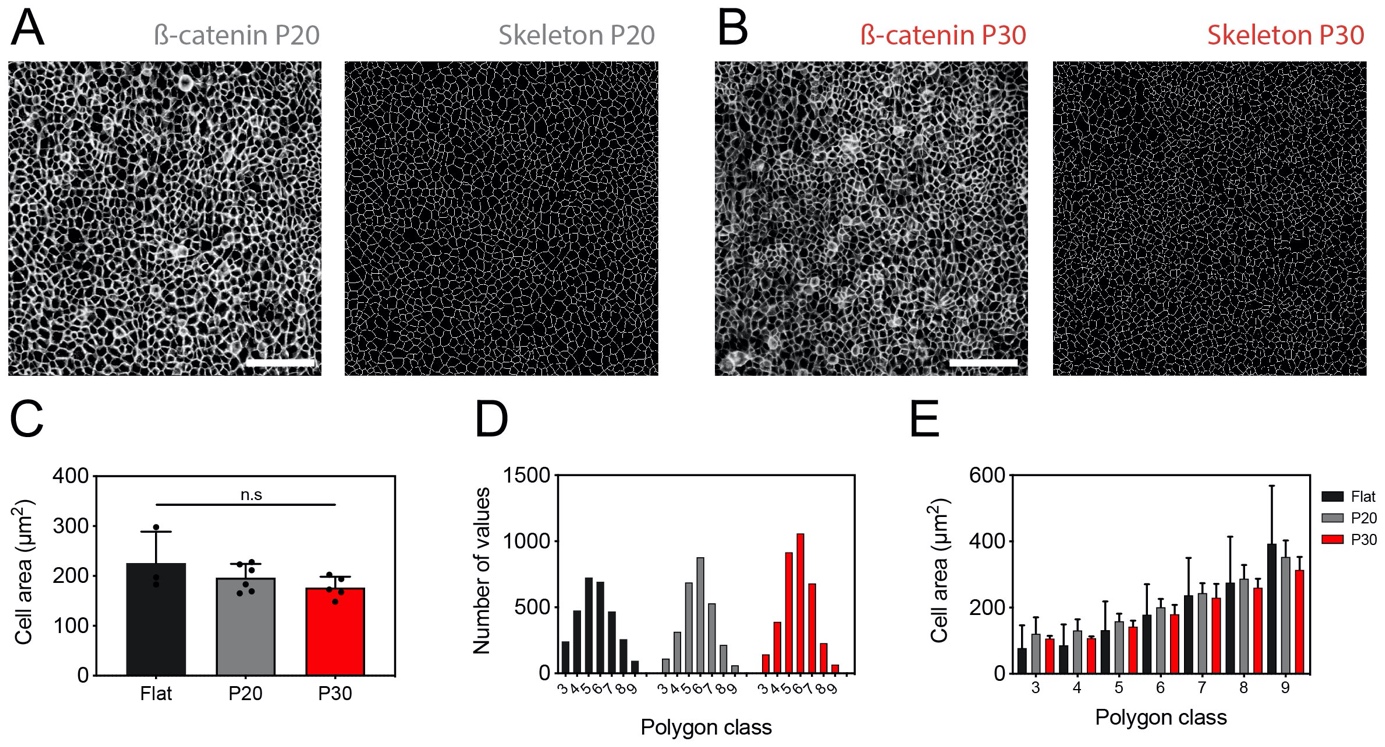

**Supplementary Figure S2 –** **Mean cell area and polygon class.** Typical maximum intensity projection image (left) and corresponding skeleton image (right) of a MDCK monolayer stained for ß-catenin and grown on (A) P20 and (B) P30 corrugated hydrogels. Scale bars correspond to 125 µm. (C) Mean cell area and (D) distribution of polygon classes of epithelial tissues grown on flat (black), P20 (grey) and P30 (red) hydrogels. n= 1100 (flat in black), 1500 (P20 in grey) and 1700 (P30 in red) obtained from 3, 6, 5 replicates respectively. (E) Mean cell area versus polygon class of epithelial monolayers grown on flat (black), P20 (grey) and P30 (red) hydrogels. n.s. is not significant.

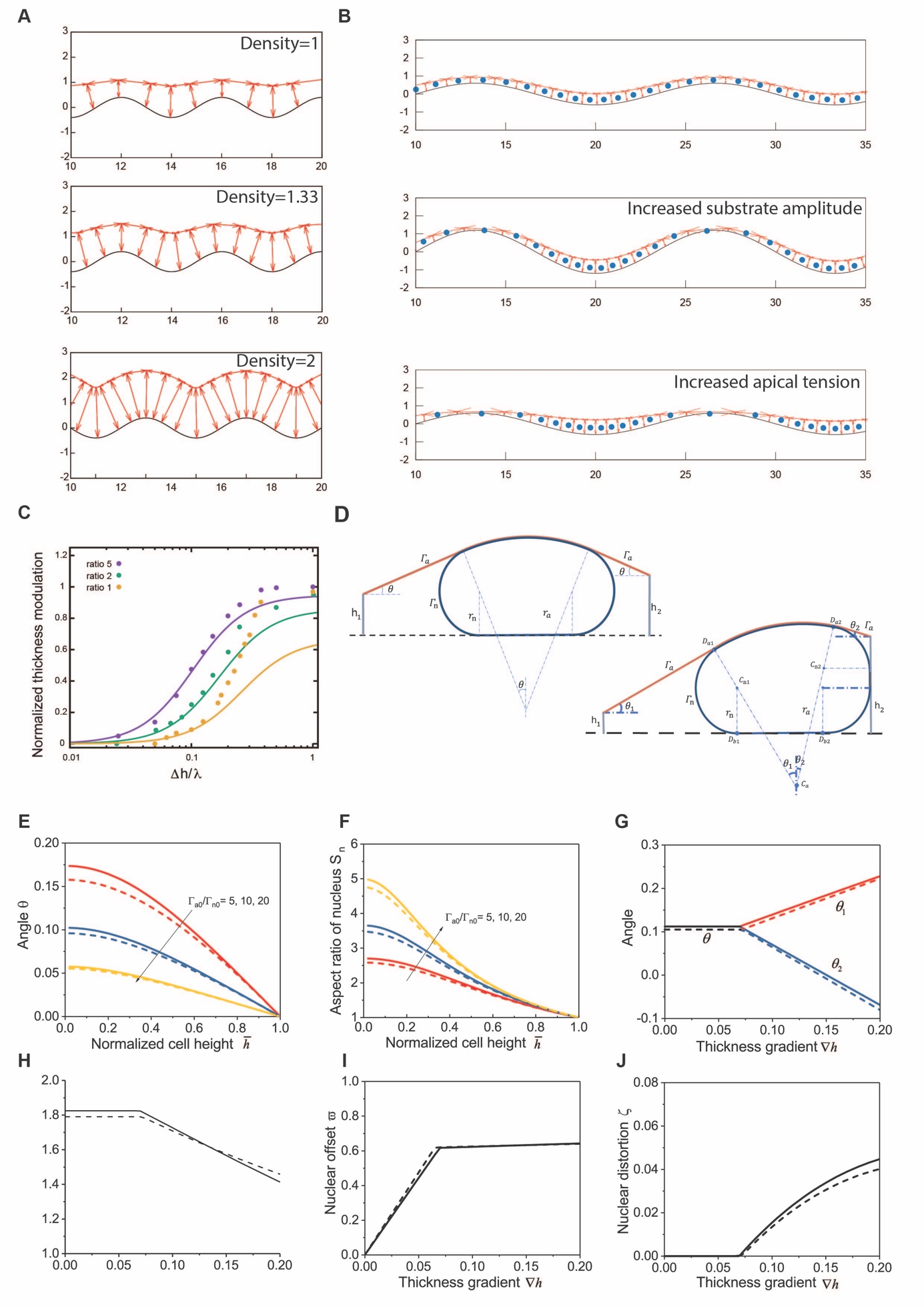

**Supplementary Figure S3 - Schematic of the model and sensitivity analysis.** (A) Simulation of the vertex model on curved substrate with increasing density (4, 6, and 8 cells per wavelength, from top to bottom), showing that increasing density decreases thickness modulations. (B) Simulation of human keratocyte on curved substrate (Mabasseri et al, see SI Theory Note for details), modelled with $\frac{\Gamma_{a}}{\Gamma_{l}}\approx5$, showing density modulations even for large wavelengths (top), which are amplified further to the point of crest/top de-wetting when doubling the substrate amplitude (middle), or when doubling apical tensions $\frac{\Gamma_{a}}{\Gamma_{l}}\approx10$ (bottom). (C) Comparison for the thickness modulation $-\Omega$ between analytical theory (thin lines) and vertex simulations (dots), for $\frac{\Gamma_{a}}{\Gamma_{l}}=5$ (purple), $\frac{\Gamma_{a}}{\Gamma_{l}}=2$ (green) and $\frac{\Gamma_{a}}{\Gamma_{l}}=1$ (purple), showing good agreement for large wavelengths $\lambda$ (normalized by average cell thickness $\Delta h$), with corrections for small wavelengths. (D) Schematics of contact mechanics model of a nucleus subjected to apical compression, before (left) or after (right) lateral contact. (E-J) Sensitivity analysis of different model parameters/observations. Both full solutions (solid line) and analytic approximations (dash line) are given (see SI Theory Note for details). We examine the influence of normalized cell height on angle $\theta$ (E) and aspect ratio $S_{n}$ (F) for the nucleus (volume ratio $\bar{\nu}=2$ and before lateral contact). We also examine the influence of local thickness gradient on angles (G), aspect ratio (H), nuclear offset compared to the cell center of mass (I), and nuclear distortion (J), with normalized side length $d=1.57$ and tension ratio $\frac{\Gamma_{a}}{\Gamma_{n0}}=5$. We find on all of these metrics a sharp transition above a critical value of thickness gradients, which occurs when the nucleus reaches lateral contact (and thus cannot increase its offset, but start adopting distorted asymmetric shape because of the asymmetric contact).

**- Supplementary Theory Note -**

In this Supplementary Theory Note, we provide details on the physical modeling for a monolayer attached to a curved substrate. The equilibrium configuration of the monolayer is obtained by using a 2D vertex model describing the apical, basal and lateral tensions of cells, and considering prescribed substrate (or basal surface). The influence of the configuration of the monolayer on nuclear position and deformation is also evaluated by a contact mechanics model, which studies the mechanical interactions between a nucleus and surrounding objects (substrate and cell surfaces). We integrate these findings in minimal simulations of 2D vertex models with nuclei.

1. **2D vertex model of monolayers on curved substrates**

The equilibrium configuration of monolayers can be obtained by summing over all cells and minimizing the global energy (Honda et al, 2004; Brezavšček et al, 2012; Hannezo et al., 2014; Okuda et al., 2015; Štorgel et al., 2016; Alt et al., 2017; Dasbiswas et al., 2018; Yang et al., 2020). Here, the monolayer on a curved substrate is described by a vertex model, where the energy of a single cell is contributed by the surface energies of its apical, basal and lateral surfaces. Hence, the total free energy of the monolayer is

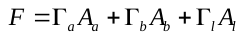
,

with
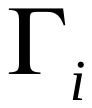
 and
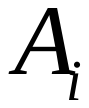
 respectively the surface tension and total area of domain *i* ( *i*=*a*, *b*, *l* respectively denotes apical, basal and lateral surface).

In this study, spatial modulation of the substrate is only patterned in one direction (along z-axis), so a 2D description of the monolayer (neglecting the in-plane direction perpendicular to the pattern) can capture the basic features of the equilibrium configuration. Moreover, confluency allows us to consider smooth continuous variations of surfaces across the monolayer. We define the spatial modulation of the basal surface as:

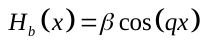
,

which is imposed on the substrate (assuming no delamination).
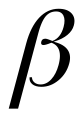
 is the amplitude and
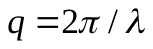
 is the wavevector of the basal (or substrate) surface. While the apical surface is free, and spatial modulation of the apical surface reads:

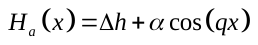
,

where
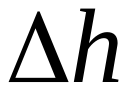
 and
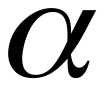
are unknowns that the monolayer can modulate to minimize its energy. Moreover, we consider the volume of the monolayer
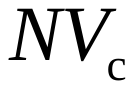
 (
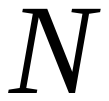
being the number of cells per wavelength or “average cell density” and
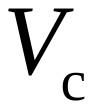
 the single cell volume) to be constant. According to
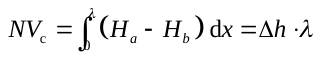
, the average thickness of the monolayer
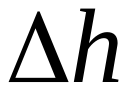
 is also constant.

We first calculate the total apical/basal areas of a monolayer over one wavelength
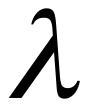
:

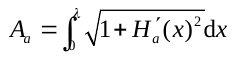
,
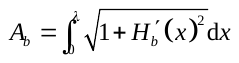
.

Considering the amplitudes of the substrate (and hence the monolayer) in experiments are much smaller than the wavelengths, we have
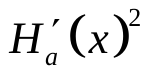
(and
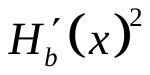
)
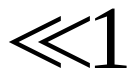
 to estimate the areas in Eq. as
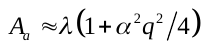
 and
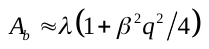
.

The total lateral area
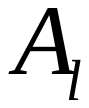
 depends on the local thickness of the monolayer (or cell height)
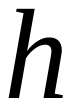
 and the cell density
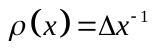
, where
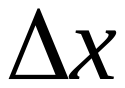
 is the length of a single cell along *x*-direction, and reads as
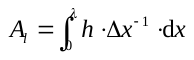
. The cell height
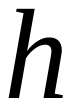
 is related to the spatial modulation of the monolayer as
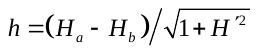
, where
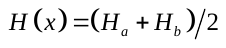
 is the mid-line of the monolayer and its slope
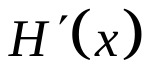
 is employed to estimate the inclination of a cell related to the *x*-axis. Considering the cell volume can be expressed as
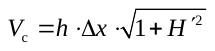
, we can rewrite the overall lateral area as

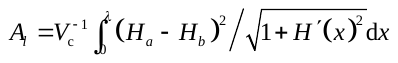
,

which can be approximated as
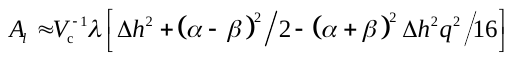
 in the limit of small amplitudes/large wavelengths (which are valid here for all wavelengths considered).

Inserting the estimates of apical, basal and lateral areas and into Eq. , the free energy of a monolayer over one wavelength is

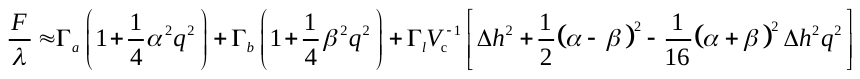
.

Minimizing this energy with respect to
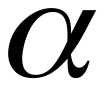
 (
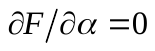
) then yields:

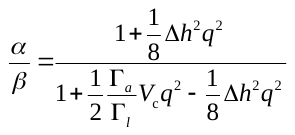
,

which clearly indicates that, for a monolayer on a curved substrate, the apical surface is generically not parallel to the basal surface (which would correspond to
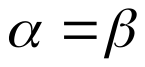
), shown as thickness modulation along *x*-axis. The dimensionless parameter
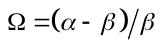
 is convenient to describe thickness modulations:

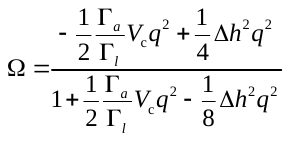
.

as the result becomes independent of the substrate amplitude
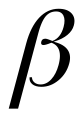
. We can see that for infinite apical tension, the thickness modulation is maximal and tends towards -1 (flat apical surface, corresponding to pure apical energy minimization), while increasing lateral tension resists this (thickness modulation approaching 0) to favor constant thickness. Intuitively, thickness modulations are also favored by small wavelengths. The analytic formula fits well with numerical results for a large wavelength
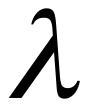
, as shown in Supp Fig. S3C. Interestingly, this suggests that for curved substrates, where the curvature changes as a function of position, curvature is generically translated into thickness/density changes. Given the fact that multiple mechano-sensitive pathways (such as YAP localization and ERK activity, as discussed in the main text) are well-known to be modulated by cell density, this minimal theory shows that the same density-sensing can be generically used to also sense curvature, without additional regulatory mechanisms/complexity. In Section 3 of this Theory Note, we compare these analytical predictions to numerical simulations of the 2D vertex model on curved substrates.

During the preparation of this manuscript, a similar criterion was put forward in a pre-print (Harmand and Hénon, 2020), although with a number of differences. In our model, we consider the average cell density *N* to be set by the seeding density and by cellular proliferation/loss (rather than from balance of tensions as assumed in Harmand and Hénon, 2020). This is expected to be valid for confluent tissues in the absence of delamination, which thus cannot adjust their overall areas at constant cell number (and given that different densities can be simply attained by changing the seeding density).

Next, we wish to understand how thickness modulation of the monolayer can influence nuclear position and morphology.

2. **Contact mechanics of nucleus**

In this section, we will evaluate the consequence of such thickness modulations for other features of the monolayer, such as nuclear shape and positioning, given the key role of the nucleus for mechano-sensation (Hamouda et al, 2020).

For a single cell or confluent cells attached to a substrate, the nucleus usually sits on the substrate and compressed by the apical surface (Versaevel et al., 2012; Iyer et al., 2020). Here, we propose a minimal contact mechanics model to evaluate the influence of the height of the apical surface (i.e. local monolayer thickness), and especially the inclination of the apical surface related to the basal surface (or substrate), due to thickness modulation, on nuclear position and deformation. For simplicity, we model the nuclear envelope as a membrane which only sustains in-plane tension. Although nuclear mechanics is much more complex (Hamouda et al, 2020), this is a minimal assumption that is sufficient to capture the tendency of the nucleus to adapt its shape and position to minimize deformation energy. The nuclear envelope is inflated by a uniform pressure
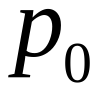
. Nucleus contacting either with the cell surfaces or the substrate is considered to be free from friction and adhesion (Fig. 3C), which we expect to be a suitable approximation for the long-time scales considered here as we only look at steady-state solutions.

Under a uniform pressure load, a membrane has spatially uniform tension and curvature. Thereafter, the composite membrane (i.e. the contact region of the apical surface and nuclear envelope) and the non-contact region of the nuclear envelope are circular arcs with radii of curvature denoted as
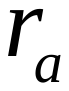
 and
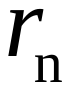
, respectively. While the non-contact region of the apical surface is a straight line, and the regions of the nucleus in contact with the rigid substrate or lateral surface are also flat. For non-adhesive contact of membranes, in-plane tensions keep the same direction across the contact edges (i.e.
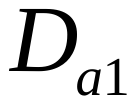
,
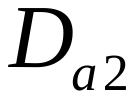
,
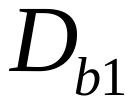
,
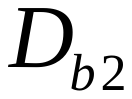
). This implies that, both the straight portion of apical surface and the non-contact region of nuclear envelope are tangent to the composite membrane, which lead to the collinearity of centers of curvature (e.g. that of the nuclear envelope
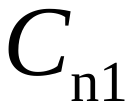
(or
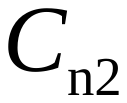
) and of the composite membrane
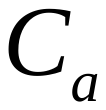
) , and the contact angle at the nucleus–substrate contact edge (
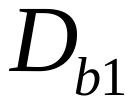
,
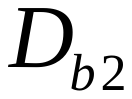
) is zero. Note that this model assumes that nucleus adapts to the shape driven by active interfacial tensions, neglecting the feedback from nuclear mechanics to overall cell shape.

a. *Nucleus without lateral contact*

We first consider a nucleus without lateral contact, as shown in Supp Fig. S3D (left panel). For a constant apical tension
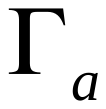
, force balance in *x*-direction requires the same angle between the straight apical surface and the substrate on two sides, denoted as
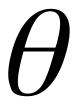
, and symmetric nuclear deformation.

For a membrane under a uniform pressure load, the in-plane tension and curvature follow the Laplace law. In the 2D model, the radii of the composite membrane (of the apical surface and the nuclear envelope) and the non-contact region of the nuclear envelope respectively yield:

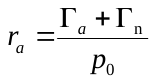
,
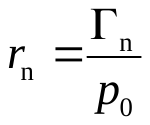
,

where
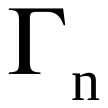
 is the tension in the nuclear envelope and will change with nuclear deformation:

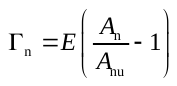
,

where
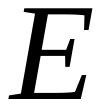
 is the area expansion modulus of the nuclear envelope,
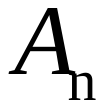
 and
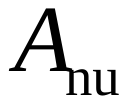
 are respectively the areas of nuclear envelope in its current and unstretched state. We have
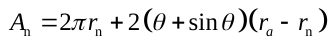
 for a symmetrically deformed nucleus. For convenience, we also consider the free state of the nucleus, which is a round membrane subjected to inner pressure
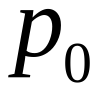
. The nuclear envelope will engender an initial tension
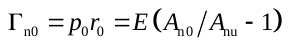
, where
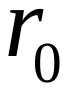
 is the radius of the nucleus in free state and initial area
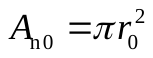
. Besides, the deformation of the nucleus should satisfy the geometric constraints

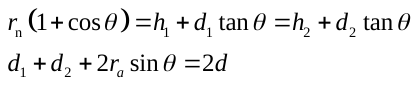
,

where
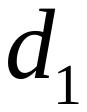
 ( or
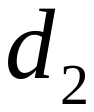
) is the distance (along the substrate surface) between contact edge
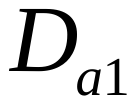
 (or
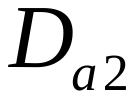
) and the left (or right) lateral boundary,
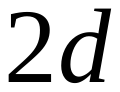
 is the total side length (along the substrate surface) of the cell. After simple manipulations of Eq. , we find that the nuclear deformation is only relevant to the average cell height
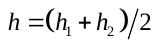
, but independent on the thickness difference
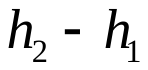
. After the normalization of geometric quantities by nuclear radius
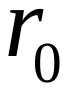
:
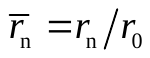
,
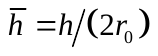
, and
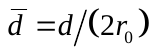
, Eqs. - can be rewritten as

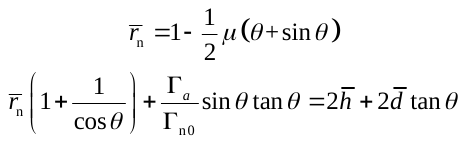
,

where
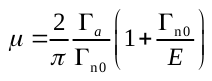
. For moderate tension and elastic deformation of the nuclear envelope, we have
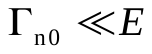
 and
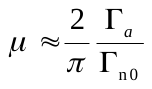
.

For small angle
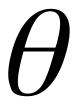
 , we can simplify Eq. and get analytic expressions of geometric variables (
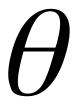
,
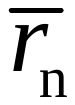
):
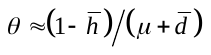
,
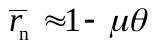
, which are close to the full solutions for the whole range of normalized cell height
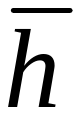
 from 0 to 1 (Supp Fig. S3E). Here, we assume constant cell volume
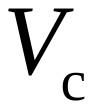
 and introduce the volume ratio
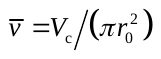
 to characterize the relative size of the nucleus compared to the whole cell. For constant cell volume, the relation between the cell height
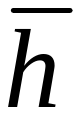
 and side length
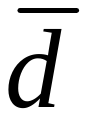
is
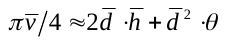
, which yields
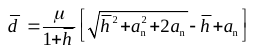
with
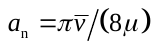
.To characterize the deformation of nucleus, we introduce the aspect ratio of nucleus
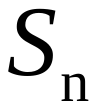
, which denotes as the length ratio of the major axis to minor axis (see Supp Fig. S3D for definitions and schematics). The aspect ratio
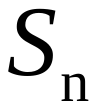
 can be well approximated by the analytic formula (Supp Fig. S3F).Under the compression of apical surface, a decrease in cell height will lead to the increase the aspect ratio and squash the nucleus flat (Supp Fig. S3F).

Strikingly, although the local thickness gradient has no impact on nuclear deformation in this case (Supp Fig. S3H, left regime), it can modulate the relative position of the nucleus inside a cell, characterized by the offset (see Supp Fig. S3D for definitions and schematics). Continuous increase of will drive the nucleus towards the portion with higher cell height (Supp Fig. S3D, G, I), until the nucleus contacts with the lateral boundary. This shows strong similarity with our experimental findings on nuclear positioning on curved substrates, and we will thus discuss it in the following.

b. *Nucleus with lateral contact*

Under lateral contact, the nuclear configuration is no longer symmetric, as shown in the right schematic of Supp Fig. S3D. Laplace law still holds, while to evaluate the tension of nuclear envelope by Eq. , we now have the nuclear surface area . The asymmetric nuclear profile yields the following geometric constraints:

,

where , andis the length of lateal contact line. Finally, the asymmetric nuclear deformation, governed by three parameters (, , ), can be obtained by solving the following equations:

.

For asymmetric nuclear deformation (i.e. the scenario with lateral contact), Eq. can be linearized by considering small angles and . We can obtain ,. The distortionis introduced to characterize the asymmetry of nuclear deformation.

As shown in Supp Fig. S3G-J, before lateral contact, local thickness gradient only promote the relocation of the nucleus (i.e. a linear increase in offset , Supp Fig. S3I), which keeps the same symmetric shape (i.e. constant aspect ratio, Supp Fig. S3H and no distortion , Supp Fig. S3J). Above a critical thickness gradient, which occurs when the nucleus reaches lateral boundary, the nucleus no longer increases its offset (Supp Fig. S3I), but sustains distorted asymmetric deformation because of the lateral contact (Supp Fig. S3J).

The specific expressions of metrics characterizing the position and deformation of nucleus, are listed in Table S1.

**Table S1** Formulas of geometric metrics in contact mechanics model

| **Variable** | **Nucleus before**  **lateral contact** | **Nucleus after lateral contact** |
| --- | --- | --- |
| Aspect ratio |  |  |
| Offset |  |  |
| Distortion |  |  |

(): length of nuclear major (minor) axis; (): distance (along the substrate surface) between centers of curvature and left (right) lateral boundary; () : height of ().

3. **Numerical simulations**

In this section, we give details on the discrete 2D vertex modeling of the monolayer on curved substrates, compare the thickness modulation shown in numerical simulations to those predicted by the vertex model with continuum assumptions in Section 1 and observed in experiments. Besides, Nuclei, treated as point particles, are incorporated into the discrete vertex model to evaluate the influence of the thickness modulation on nuclear positioning.

a. *Vertex model of monolayer*

To simulate the 2D vertex apico-basal configuration of cells on curved substrate, we assume that each cell *i* is described by the basal vertices $b_{i}$, $b_{i+1}$ and apical vertices $a_{i}$, $a_{i+1}$ (altogether ascribing cell area $A$), so that the single cell energy reads:

$E_{i}= \Gamma_{a} |a_{i}-a_{i+1}|+\Gamma_{l}(\left| a_{i}-b_{i} \right|+\left| a_{i+1}-b_{i+1} \right|)+K\left( A-A_{0} \right)^{2}$,

where $\Gamma_{a}$, $\Gamma_{l}$ are, as above, apical and lateral tensions (basal tensions vanish from the model as discussed above), and $K$ the compressibility modulus which we take very large to ensure cell area remains constant (in the simulations, we set $\Gamma_{l}=1, A_{0}=1$ without loss of generality, use $K=20$, and explore changes in $\Gamma_{a}$). Absolute values denote Euclidian distance between two vertices, so that $\left| a_{i}-b_{i} \right|$represents the height of the lateral area shared by cells *i* and *i*-1. We set the cell density by the number of cells in a given wavelength upon initialization, and maintain the first and last cell fixed (confluency condition). In all cases apart from when we explore changes in density, we set initial cellular width to 1 (so that average thickness is also 1 given $A_{0}=1)$. We minimize this energy using a Monte-Carlo algorithm with very low temperature, randomly changing the positions of apical and basal vertices by $(\delta x, \delta z)$ (each component out of a uniform distribution $[0,0.025])$ at each step, and for long times to ensure the energy has converged. When we change at random the position of an apical vertex, both $\delta x$ and $\delta z$ components are picked at random, whereas for a basal vertex, the $\delta x$ component is also picked at random, while $\delta z$ is then imposed by substrate curvature equation $z=\beta cos(2\pi x/\lambda)$, to ensure that the basal surfaces remain attached to the substrate. These simulations show excellent agreements with the analytical criterion derived in Section 1 for large wavelength (Fig. 2E-G, S4C), although the theory underestimates thickness modulation for small wavelengths (at which point the continuum theory breaks down, and strong localization of vertices in valley/crests occur, notably for integer ratios of wavelengths over number of cells (Fig. 3G).

b. *Parameter measurements, fitting and comparison to data*

All parameters in the theory/simulations (overall cellular density, cell height, substrate amplitude/wavelengths) can be measured, as discussed in the main text, except $\Gamma_{a}/\Gamma_{l}$, which need to be fitted. For this, as shown in Fig. 2A-D, we plotted local thickness as a function of local substrate amplitude $z=\beta cos(2\pi x/\lambda)$. Theoretically, this is predicted to be linear, with the slope being exactly $\Omega$, and the intercept at $z=0$ giving the average cell height. This was well-verified in the data (Fig. 2B, D), providing a good test of the model, and allowing us to extract the free parameter $\Gamma_{a}/\Gamma_{l}$.

However, both P20 and P30 are in the small wavelength regime and show thickness modulation already saturated close to 1, so that we can only set a lower bound on $\Gamma_{a}/\Gamma_{l}$, finding that the thickness modulation we observe is consistent with $\frac{\Gamma_{a}}{\Gamma_{l}}>1$. Interestingly, increasing cellular density has a similar effect as increasing lateral tension (as increasing cellular density creates more lateral surfaces in a given unit length), i.e. decreasing thickness modulations, as shown in Supp Fig. S3A. This mirrors findings of YAP localization at high densities (Fig. 4A, E) where curvature sensing is abolished.

Furthermore, Mabasseri et al. found that in human keratocytes, 2.5-fold lower cell densities were observed in crests vs valleys (Mobasseri et al., 2019). From their image, we estimated wavelengths $\lambda\approx300\mu m$ (corresponding to around 15 cell lengths) and substrate amplitudes $\beta\approx15\mu m$. We find that 2.5-fold lower densities in crests are well-captured by setting $\frac{\Gamma_{a}}{\Gamma_{l}}\approx5$ (Supp Fig. S3B, top panel), arguing for applicability of the theory in other settings, and that as YAP nuclear localization has been shown to be density-dependent (see discussion in main text), such theory gives a simple explanation for curvature-sensing, by converting substrate curvature into density modulations. Furthermore, Mabasseri et al. found that increasing substrate amplitude (roughly two-fold) caused the epithelium to de-wet from the crests (Mobasseri et al., 2019). In our model, this could be reproduced, as for $\beta>\Delta h$, large enough density/thickness modulation are enough to make thickness at crests to approach zero. This experimental finding could thus be well-reproduced by doubling the substrate amplitude $\beta$ in the simulations (Supp Fig. S3B)

c. *Vertex model incorporating nuclear positioning*

To integrate the findings on nuclear positioning in simulations, we thus ran our 2D vertex model, adding nucleus as point particles maintained onto the center of mass of a cell by a spring force, and subject to an additional force proportional to the difference in its two lateral heights (i.e. the regime of linear response to thickness gradients shown analytically in Table S1 and Supp Fig. S3I). Finally, forces acting on nuclear position $n_{i}$ are

$f_{i}= {-K}_{n}\left( n_{i}-n_{0i} \right)-f_{r}\left( \left| a_{i}-b_{i} \right|-\left| a_{i+1}-b_{i+1} \right| \right)$,

where equilibrium position $n_{0i}$ of the nucleus is calculated as the center of mass of cell *i* (we assume the nucleus is actively maintained at this position and model this constrain as a spring force, setting $K_{n}=1$ without loss of generality), and $f_{r}$ quantifies the strength of the forces from thickness gradients $(\left| a_{i}-b_{i} \right|-\left| a_{i+1}-b_{i+1} \right|)$ on setting nuclear position/offset from the center.

Nuclei are thus slave to the 2D vertex model under this assumption. Interestingly, we find that nuclei on the side of the substrate patterns (where gradients in thickness are the largest) tend to move towards valleys, as observed in data (Fig. 3H-J). Strikingly, this effect becomes highly amplified for very small wavelengths (only a few cells per wavelength, at which the continuum approximation in Section 1 is expected to break down). As shown in Fig. 3I and discussed in main text, we observe a strong quantization of the response in particular for 2 cells per wavelength, as the energy minimum is obtained when a pair of cells each occupies a half pattern, with lateral surfaces sitting at the crests/valleys (Fig. 3G). This maximizes thickness gradients and causes all nuclei to be localized in valleys (in strong similarity to the data for 30-micron wavelengths, Fig. 3I, J). This does not occur when the cell/wavelength ratio is not an integer, as equilibrium configurations are disordered with respect to the periodic substrate (Fig. 3H, J).
